# Unsupervised, piecewise linear decoding enables an accurate prediction of muscle activity in a multi-task brain computer interface

**DOI:** 10.1101/2024.09.09.612102

**Authors:** Xuan Ma, Fabio Rizzoglio, Kevin L. Bodkin, Lee E. Miller

**Affiliations:** Department of Neuroscience, Northwestern University, Chicago, IL, United States of America; Department of Neurobiology, Northwestern University, Evanston, IL, United States of America; Department of Biomedical Engineering, Northwestern University, Evanston, IL, United States of America; Shirley Ryan AbilityLab, Chicago, IL, United States of America; Department of Physical Medicine and Rehabilitation, Northwestern University, Chicago, IL, United States of America

**Keywords:** intracortical BCI, multi-task, neural decoding, EMG, piecewise linear

## Abstract

**Objective:** Creating an intracortical brain-computer interface (iBCI) capable of seamless transitions between tasks and contexts would greatly enhance user experience. However, the nonlinearity in neural activity presents challenges to computing a global iBCI decoder. We aimed to develop a method that differs from a globally optimized decoder to address this issue.

**Approach:** We devised an unsupervised approach that relies on the structure of a low-dimensional neural manifold to implement a piecewise linear decoder. We created a distinctive dataset in which monkeys performed a diverse set of tasks, some trained, others innate, while we recorded neural signals from the motor cortex (M1) and electromyographs (EMGs) from upper limb muscles. We used both linear and nonlinear dimensionality reduction techniques to discover neural manifolds and applied unsupervised algorithms to identify clusters within those spaces. Finally, we fit a linear decoder of EMG for each cluster. A specific decoder was activated corresponding to the cluster each new neural data point belonged to.

**Main results:** We found clusters in the neural manifolds corresponding with the different tasks or task sub-phases. The performance of piecewise decoding improved as the number of clusters increased and plateaued gradually. With only two clusters it already outperformed a global linear decoder, and unexpectedly, it outperformed even a global recurrent neural network (RNN) decoder with 10-12 clusters.

**Significance:** This study introduced a computationally lightweight solution for creating iBCI decoders that can function effectively across a broad range of tasks. EMG decoding is particularly challenging, as muscle activity is used, under varying contexts, to control interaction forces and limb stiffness, as well as motion. The results suggest that a piecewise linear decoder can provide a good approximation to the nonlinearity between neural activity and motor outputs, a result of our increased understanding of the structure of neural manifolds in motor cortex.

## Introduction

An intracortical brain-computer interface (iBCI), which converts the spiking activity of simultaneously recorded motor cortical neurons into user intentions, is a promising tool for assisting individuals suffering from paralysis or other neuromotor disabilities [1–4]. Key to this function is the “decoder”, an algorithm that is trained to find a reliable mapping between neural activity and motor outputs (Figure 1A). An iBCI decoder is generally built using ordinary supervised machine learning algorithms, an approach that has been demonstrated quite successfully for applications involving a limited range of motor behaviors. Most existing motor iBCIs are kinematic in nature, allowing their users to control computer cursors [5,6] and robotic limbs [2,7,8]; a few provide simple force control [9]. However, an iBCI that seeks to restore a more natural interaction with objects must be able to control kinematics and forces together. This is where current iBCIs fall short, as kinematic decoders require an unnatural strategy to exert forces, for example, at the time of contact with an object during the onset of grasp [10]. A potential solution could be an iBCI that directly controls muscle activity, mimicking the control mechanism of our neuromuscular system.

**Figure 1.**
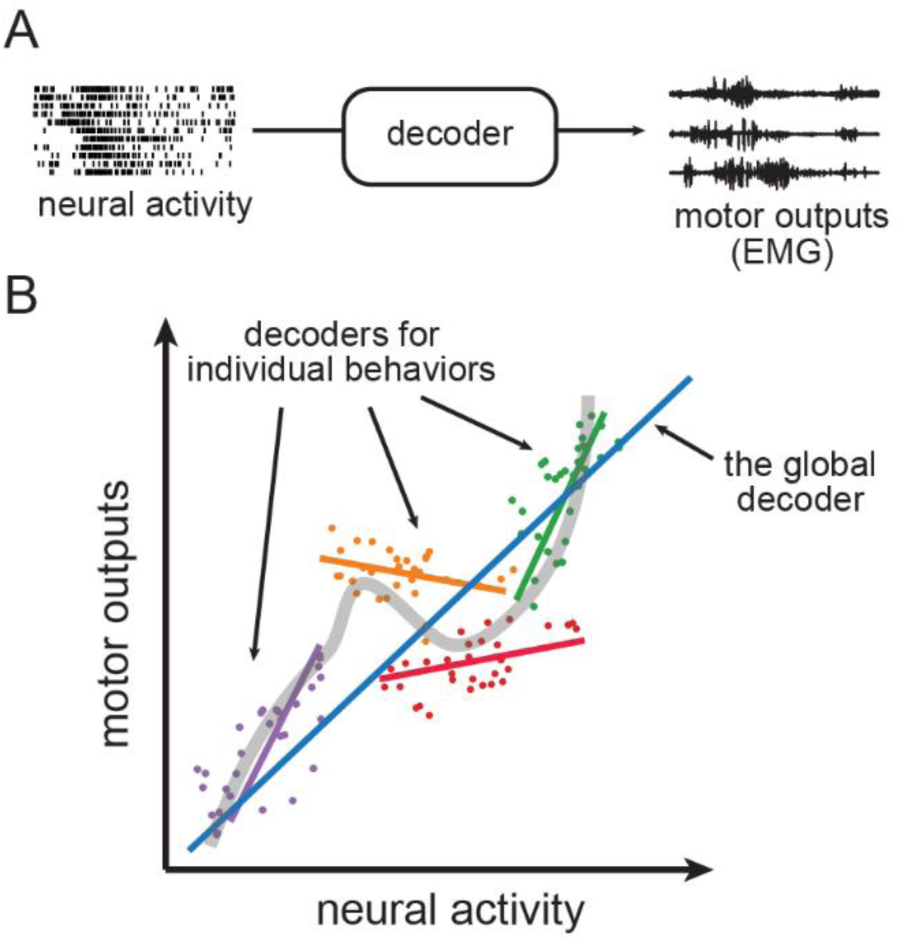
The nonlinearity in neural activity makes it difficult to fit a global decoder across multiple tasks distributed across a low-dimensional neural manifold. **(A)** A decoder maps neural activity to motor outputs. **(B)** Fitting the relationship between neural activity and motor outputs. The dots of different colors represent the data samples from different hypothetical tasks, while the colored lines denote linear decoders fit to those samples. The blue line represents a global linear decoder fit to all the data samples. The gray curved line shows a nonlinear fit over all data samples from different tasks.

**Figure 2.**
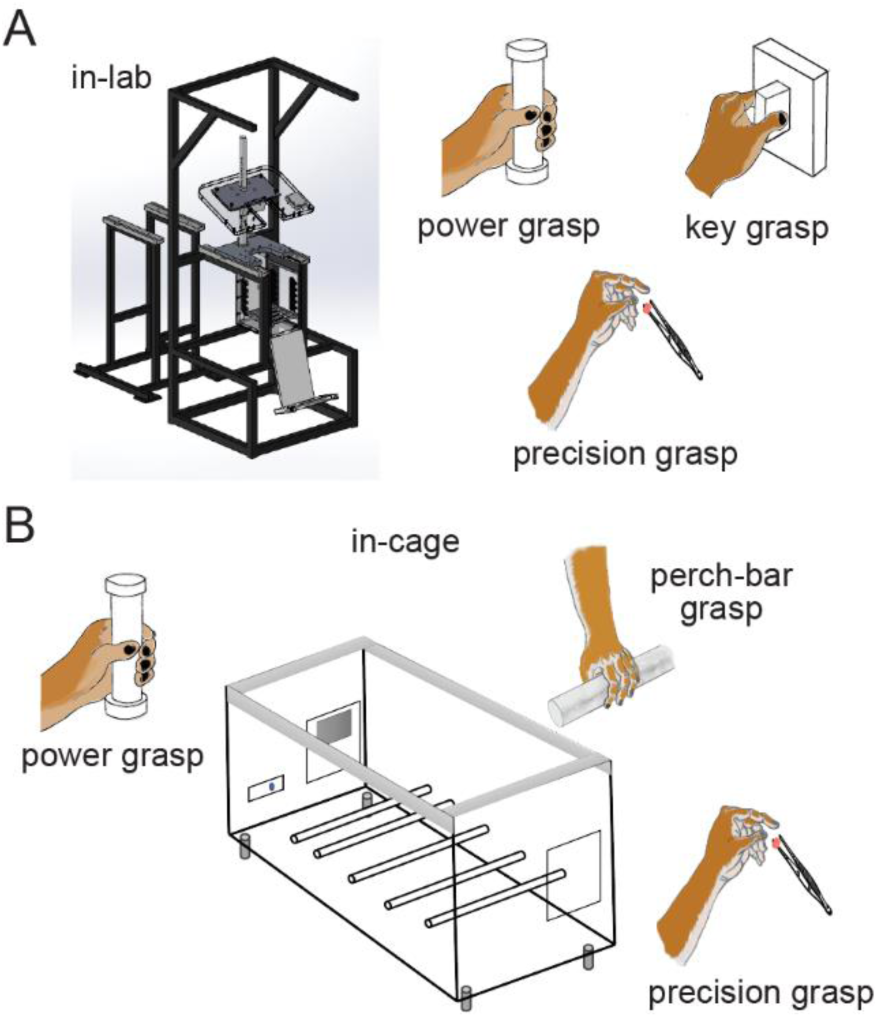
The environments where the tasks took place. **(A)** The in-lab environment. Monkeys were seated in a standard primate chair, and were trained to perform power, key, and precision grasp tasks. **(B)** The in-cage environment. Monkeys were put inside a large plastic cage. Perch bars and instrumented devices were mounted inside the cage. Monkeys were trained to perform power and precision grasps, similar to those in the lab, and performed a series of grasps on the perch bars when they walked between ends of the cage.

Our group pioneered a Wiener filter-based decoder of muscle activity, rather than movement kinematics [11]. We predicted the electromyographs (EMGs) from neural signals in motor cortex (M1) with R^2^ values ranging from 0.5 to 0.9. When used to modulate the intensity of functional electrical stimulation (FES), these real-time EMG predictions allowed the monkeys to perform the same task despite paralysis of hand muscles induced by a transient peripheral nerve block [12]. However, in later experiments, neither a Wiener filter-based or long short-term memory (LSTM) network-based EMG decoders generalized across a set of wrist tasks that required dynamical loads ranging from isometric contractions to unloaded movements [13]. An analogous observation was made for a decoder of reach velocity in a precision reaching task; following training on the initial, primary phase of movement, the decoder generated poor predictions of velocities during a subsequent corrective phase [14].

The effectiveness of linear decoders for individual tasks or sub-movements, as illustrated by the toy data samples and linear fits in Figure 1B, aligns with the linear relationship reported between M1 neural activity and motor outputs by previous studies [15–17]. However, since the motor outputs required for individual tasks usually fall within a limited range, these linear relationships tend to be local and do not capture the nonlinearities present across a broader range of motor behaviors [18–20]. This poses challenges in developing an iBCI system capable of handling multiple tasks.

Rather than developing a highly generalizable algorithm capable of predicting motor outputs for new tasks, a more feasible solution may be to train the decoder with data from a variety of tasks, allowing it to capture the “global” characteristics of the neural activity.

As the number of tasks increases, the accuracy of the predictions for any given task by a “global” linear decoder will likely decrease. It will “miss” some tasks and tend to perform worse than the decoders trained individually (Figure 1B). When we applied this approach to the wrist tasks described above, the linear decoder predictions improved but fell well short of the single-task decoders. In that study, the LSTM decoder improved predictions substantially [13], but it remained uncertain whether a global nonlinear decoder could achieve satisfactory performance when applied to a larger, more diverse range of tasks.

The performance and efficiency of these networks is significantly influenced by the selection of hyperparameters and the available data. LSTMs in particular are notoriously slow to train on larger datasets. Recently, with large language models gaining prominence [21], LSTMs have been replaced by transformer-based models, which are faster to train and can more effectively make use of vast amounts of neural recordings [22,23]. The expectation is that such pretrained large models could accommodate the variations in neural activity across time, tasks, and subjects for iBCI applications. However, the consumption of computing resources is still substantial: for instance, POYO-1 required 8 Nvidia A40 GPUs for training over 5 days, and additional fine-tuning was necessary before it could be used [23].

In this study, rather than pursuing a globally optimal linear or nonlinear decoder, we exploited the local linearity illustrated in Figure 1B. We used linear and nonlinear dimensionality reduction techniques to discover the underlying neural manifolds [24] and applied unsupervised algorithms to identify clusters within those spaces. We computed a series of piecewise linear decoders of arm and hand muscle activity, each accurate within a restricted region of the low-dimensional neural manifold. We activated specific decoders according to the cluster each new neural data point belonged to. We tested this piecewise decoding using a dataset comprising a diverse set of behaviors, some that were trained and others that were innate. With only two clusters, our approach already outperformed a global linear decoder; unexpectedly, it outperformed even a global LSTM decoder when the number of clusters reached 10-12. These findings suggest that piecewise linear decoding can achieve a good approximation to the nonlinearity between neural activity and motor outputs and provide a lightweight solution for multi-task iBCI applications.

## Methods

### Subjects and behavioral tasks

Two adult male rhesus monkeys (Macaca mulatta), T and P, were used in this study. Both were trained to perform a series of behavioral tasks in two distinct environments (Fig. 1B). The first we termed the “in-lab” environment; the monkeys were seated in a standard primate chair with the body posture fixed and one arm restrained as they performed three types of tasks. Two of the tasks required the monkeys to reach and grasp an instrumented device placed in front of them with their free hand. The device was either a 1.5cm diameter cylinder facilitating a power grasp with the palm and the fingers, or a small rectangular cuboid (1 x 2 x 2 cm) facilitating a key grasp with the thumb and the lateral of the index finger. Force sensitive resistors (FSRs) were attached on opposite sides of each device, which controlled the movement of a cursor on a monitor in front of the monkey. The sum and the difference of the forces controlled the position on the vertical and horizontal axes, respectively. The monkey had to keep his hand resting on a touch pad for a random time (0.5 – 1.0 s) to begin each trial. After the hold, one of three rectangular force targets appeared on the screen together with an auditory go cue. The monkey was required to place the cursor into the target and hold for 1 s in order to receive a fluid reward. The third task required the monkeys to grasp small food pellets held with tweezers at arm’s length by the experimenter. The monkeys typically used a precision grasp involving the thumb and the tip of the index finger, but this action was less constrained than the other two. For the power grasp and key grasp tasks, we detected the onset of grasp based on the measured forces, designating the period from 0.5 s before to 1 s after the force onset time as a trial. For the food pellet retrieval, or “precision grasp” task, we monitored the monkeys’ movements with industrial cameras (Ximea Inc., Münster, Germany). The frames in the video were synchronized to the neural recordings using pulses generated by the Cerebus system. We then identified the moment when the monkeys touched the pellet based on those videos, again using the period 0.5 s before and 1 s after as a trial.

The second “in-cage” environment, was a plastic cage measuring 2 x 1 x 1 meters, allowing the monkeys to move freely without constraints. We placed a single monkey in the cage at a time and recorded wirelessly from both primary motor cortex and a variety of muscles in the arm contralateral to the array (details below). The monkeys’ behavior was monitored using the same video recording system as in the lab. At one end of the cage, we mounted a duplicate of the in-lab power-grasp device, actuated like the in-lab device, but with the monkey required to balance while squatting on the floor of the cage, and to initiate the task by pushing a button instead of the touch pad. At the opposite end, a small opening allowed the monkeys to stand and interact with the experimenter, such as grasping small food pellets, an in-cage version of the precision grasp task. Five rods spanning the width of the cage and positioned 10 cm above the floor served as perch bars as the monkeys moved between the power and precision grasp ends of the cage. While walking back and forth along the length of the cage, the monkeys typically performed a series of power grasps on these bars. Given the difficulty in controlling the timing of movements in such an unconstrained setting, we relied again on video recordings to identify and time-align these grasps. Together with the lab tasks, our analyses focused on these three in-cage tasks: discrete power and precision grasps at either end of the cage, and the locomotion-related grasps on the perch bars. For the power grasp task, we obtained the force onset time in the same way as for the in-lab power grasp. For the other two tasks, we identified the moments when the monkey touched the food pellet or placed the palm on the perch bars based on video recordings to extract 1.5-s trials as the lab. In addition to these defined cage tasks, the monkeys engaged in various spontaneous behavior as well.

We began each experimental session with the in-lab power and key grasp tasks in interleaved trials, followed by a block of the in-lab precision grasp task. Typically, these sessions lasted 35 to 45 minutes. Once finished, we removed the monkey from the primate chair and put him in the plastic cage. In-cage recordings typically began within 15 minutes after the in-lab recordings and lasted for 60 – 90 minutes.

### Surgical procedures and data collection

We implanted a 96-channel Utah electrode array (Blackrock Neurotech, Inc.) between the arm and hand representations in the so-called “PDC zone” of M1 [25] of each monkey (monkey T: left M1, monkey P: right M1). The stereotaxic coordinates of the implant site were pre-planned and finally determined during the surgery with reference to the sulcal patterns and the muscle contractions evoked by intraoperative surface cortical stimulation. We also implanted intramuscular leads in upper arm, forearm and hand muscles of the arm contralateral to the implanted array in a separate surgical procedure (monkey T: right arm, monkey P: left arm). Electrode locations were verified during surgery by stimulating each lead.

M1 activity was recorded during task performance using a Cerebus system (Blackrock Neurotech, Inc.). For the in-lab recordings, neural signals were amplified by a Cereplex-E headstage. For the in-cage recordings, neural signals were amplified, digitized at 30 kHz, and transmitted by a Cereplex-W wireless headstage. In both environments, the signals on each channel were then bandpass filtered (250 - 5000 Hz) and converted to spike times based on threshold crossings. The threshold was set with respect to the root-mean square (RMS) activity on each channel (monkey T: −5.25 x RMS; monkey P: −4.75 x RMS). The time stamp and a 1.6 ms snippet of the waveform surrounding each threshold crossing were recorded. For all analyses in this study, we used multiunit threshold crossings instead of discriminating only well-isolated single units. We applied a Gaussian kernel (S.D. = 50 ms) to the spike counts in 25 ms, non-overlapping bins to obtain a smoothed estimate of firing rate as function of time for each channel.

The EMG signals were amplified, band-pass filtered (4-pole, 50 - 500 Hz) and sampled at 2000 Hz using a 32-channel amplifier chip (RHD 2132, Intan Inc., Los Angeles, CA) and a wireless transmitter (RCB-W24A, DSP Wireless Inc., Haverhill, MA) for both in-lab and in-cage environments. In the cage, this device was placed in a backpack on the monkey’s jacket. Since the amplifier recorded monopolar EMG inputs, software differencing was performed between the paired channels after digitalization. The EMGs were then digitally rectified, low-pass filtered (4-pole, 10 Hz, Butterworth) and subsampled to 40 Hz to match the time resolution of the binned neural data. To remove occasional artifacts, the data points of each channel were clipped to be no larger than the mean plus 6 times the S.D. of that channel. Within each recording session, we removed the baseline of each EMG channel by subtracting the 2nd percentile of the amplitudes and normalized each channel to the 90th percentile across all tasks. For each monkey we recorded EMG signals from 16 muscles (Table 1).

**Table 1.** The list of muscles with intramuscular electrodes implanted*.

| Monkey | Upper arm muscles | Forearm muscles | Hand muscles |
| --- | --- | --- | --- |
| T | TRI, BI | ECU, FCU, PT, FDS, FDP, ECR, FCR, EDC, SUP | Lum, FPB, APB, 1DI, 3DI |
| P | TRI, BI | ECU, FCU, PT, FDS, FDP, ECR, FCR, EDC, EPL | Lum, FPB, APB, 1DI, 4DI |

\* Abbreviations for the muscles: ECU (extensor carpi ulnaris), FCU (flexor carpi ulnaris), ECR (extensor carpi radialis), EDC (extensor digitorum communis), FCR (flexor carpi radialis), FDP (flexor digitorum profundus), ECR (extensor carpi radialis), FCR (flexor carpi radialis), 1DI (first dorsal interosseous), FPB (flexor pollicis brevis), PT (pronator), SUP (supinator), APB (abductor pollicis brevis), Lum (the first lumbrical), FDS (flexor digitorum superficialis), 4DI (fourth dorsal interosseous), EPL (extensor pollicis longus), TRI (triceps), BI (biceps)

We conducted two styles of analysis with the neural and EMG data. In one approach, we split the data, including both in-lab and in-cage recordings, into trials as described above. In the other, focusing only on the in-cage recordings, we retained the continuously acquired data without dividing it into trials.

All surgical and experimental procedures were approved by the Institutional Animal Care and Use Committee (IACUC) of Northwestern University under protocol #IS00000367 and are consistent with the Guide for the Care and Use of Laboratory Animals.

### Piecewise linear decoders

The construction of the piecewise linear decoders is shown in Fig. 3A. In the first step, we applied either Principal Component Analysis (PCA) or Uniform Manifold Approximation and Projection (UMAP) over the full-dimensional neural recordings of the training set to find the low-dimensional embeddings of the neural activity in the latent space. PCA finds a linear manifold, as it transforms the original data into a new coordinate system such that each successive orthogonal component captures a smaller amount of variance in the data. We included the first 12 components in the subsequent analyses, which accounted for more than 70% of variance in the concatenated trials from all tasks. In contrast, UMAP finds a nonlinear manifold for neural activity by constructing a graph-based representation of the high-dimensional data, then optimizing the layout of the representation in the low-dimensional space by minimizing the cross-entropy between the two representations [26]. We used parametric UMAP [27], which learns the relationship between the original data and low-dimensional embeddings by using a neural network to minimize the same objective function as regular UMAP. This network allowed us to project new data into the low-D space without having to retrain the model, which would have been necessary with standard UMAP. For consistency with PCA, we also used the first 12 components for all subsequent UMAP-based analysis. UMAP includes a “min_dist” parameter that sets the minimum distance that adjacent data points can have in the low-dimensional representation. Smaller “min_dist” values encourage UMAP to preserve finer topological structure while larger ones preserve the broader topological structure. We set min_dist to 0.75 to emphasize broader manifold structure. An additional “n_neighbors” parameter constrains the size of the local neighborhood UMAP considers when learning the embeddings. Low n_neighbors values emphasize local structure while large ones increase the neighborhood size surrounding each point. Due to the large size of our datasets, we set n_neighbors to 1/1000 of the number training sample of datapoints to balance the tradeoff between global and local structure and minimize the computational complexity which increases with n_neighbors.

**Figure 3.**
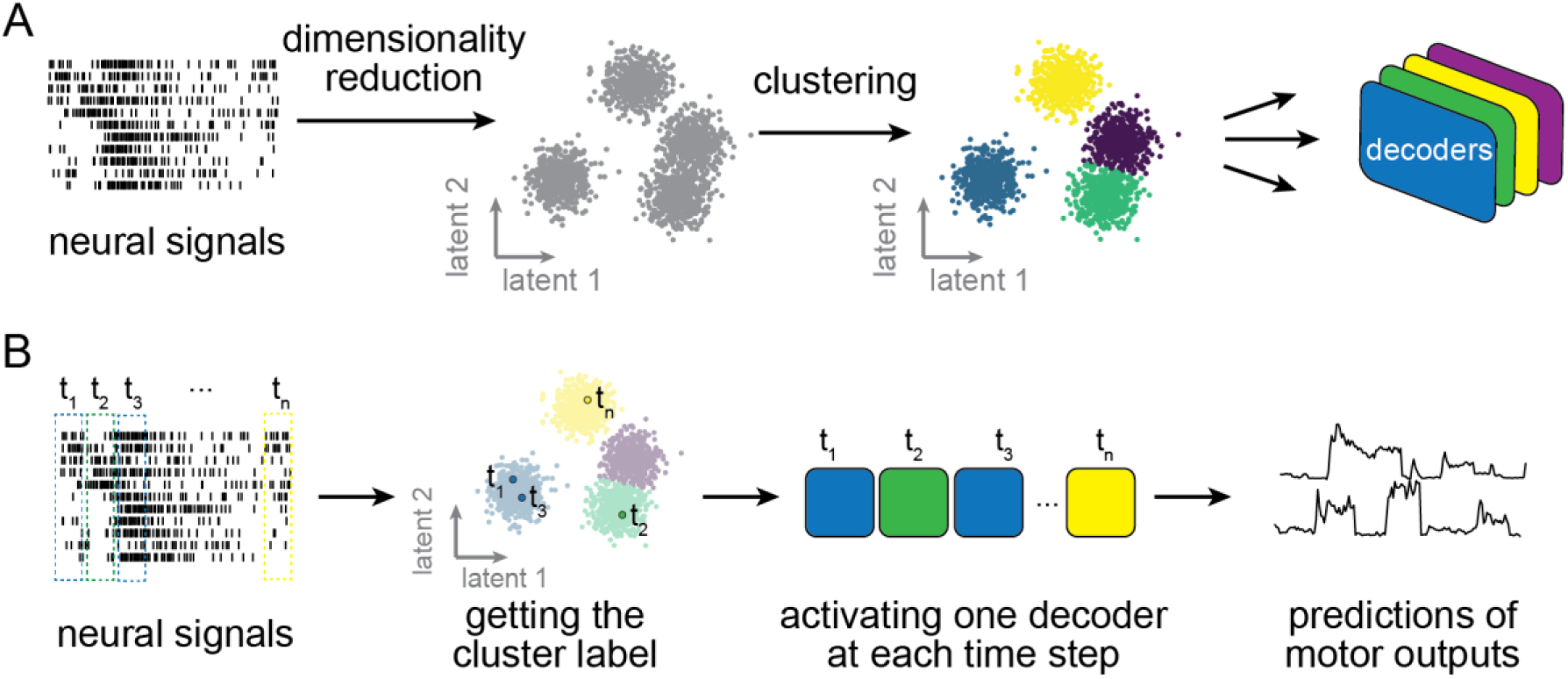
The construction of piecewise linear decoders. **(A)** The decoder training process. Full dimensional neural signals were projected into low dimensional latent space, where clusters were identified. The Gaussian mixture model (GMM) was used to identify these clusters. For each cluster (indicated by different colors), a decoder was trained using the data samples in this cluster. **(B)** The use of the trained decoders on incoming new data samples. A cluster label for the data sample at a given time *t* was obtained after passing the sample through the trained dimensionality reduction and GMM. One of the trained decoders was activated based on the cluster label to generate a prediction of motor outputs at this time step.

Following dimensionality reduction with either PCA or UMAP, we used Gaussian mixture models (GMMs) to find clusters in the neural manifolds in an unsupervised manner. GMM assumes the data points were generated from a mixture of a specified number of Gaussian distributions, with unknown mean and variance. After initially estimating the center of each distribution using a k-means clustering algorithm, we ran the expectation-maximization (EM) algorithm for 100 iterations to refine the model parameters. We set the components’ covariance to “full”, allowing them to take any shape and orientation. We used the “GaussianMixture” class in “scikit-learn” Python package [28] to implement the algorithms.

We then trained a Wiener filter-based decoder for each of the identified clusters as in several previous studies [13,29]. The filter was fitted using linear regression to predict the EMG at time *t* given neural responses from time *t* to time *t* - T, where we set T = 8 (200 ms) for all decoders used in this study:

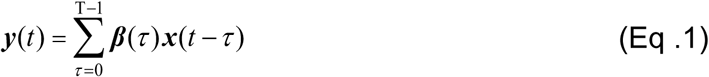

Here, ***y***(*t*) is a 16-dimensional vector of EMGs to be predicted at time *t*, while ***x***(*t*) is a 96-dimensional vector for of neural firing rates at time *t* and ***β***(*τ*) is a matrix corresponding to the filter parameters for time step *τ*. We can write the equation above in matrix form:

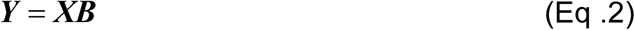

where ***Y*** is an *M*×16 matrix for the EMGs to be predicted, with *M* being the number of samples. ***X*** is an *M*×(8×96) matrix and ***B*** is an (8×96)×16 matrix of the regression coefficients to be estimated. We also added a bias term for both ***X*** and ***B***. We computed ***B*** by a ridge regression estimator:

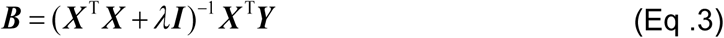

We chose ridge regression to limit the risk of decoder overfitting by penalizing solutions with large regression coefficients with the regularization term *λ*. Its value was chosen by sweeping a range of 20 values between 10 and 10^5^ on a logarithmic scale.

After training all models and decoders, new data samples were processed sequentially as illustrated in Figure 3B. At each time step, one of the decoders was activated based on proximity to the clusters identified by the GMM.

### Global decoders

We compared the proposed piecewise linear decoders to both linear and nonlinear global decoders. The global linear decoder followed the same form as described in Equation (1), with the distinction that it was trained using the data from all tasks across the two environments. Likewise, the global nonlinear decoder was a recurrent neural network with LSTM cells trained on all tasks. The network comprised one LSTM layer with 100 units and one feedforward layer with 16 outputs, to match the number of EMG channels. We used PyTorch [30] to implement this LSTM-based decoder. We tuned the hyper-parameters listed in Table 1 and selected the set enabling the best results (Table 2).

**Table 2.**
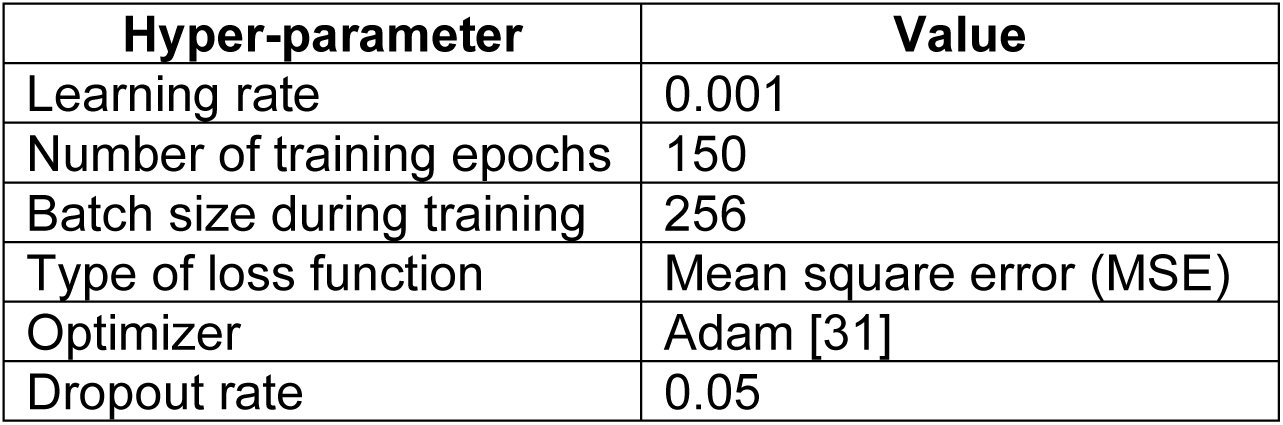
The hyper-parameters of the LSTM-based decoder.

### Evaluating the quality of clusters

To assess the quality of the clusters, we computed the “silhouette score” [32], which quantifies the proximity of a data point to other points in its own cluster compared to all other clusters. The silhouette score ranges from −1 to 1, with values closer to 1 indicating more distinct clustering and values near 0 suggesting overlaps between clusters. We computed the silhouette score for each sample, then averaged across all samples to report a single number. Assuming *x* is a data sample in cluster *C_X_*, we computed the mean distance, *d* between *x* and all other samples in *C_X_* as:

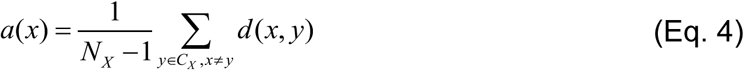

where *N_X_* is the total number of samples in *C_X_*, and *y* represents another data sample in *C_X_*. We then computed the mean distance between *x* and all the samples in some other cluster *C_Z_*, and found the mean distance from *x* to all samples in its nearest neighboring cluster as:

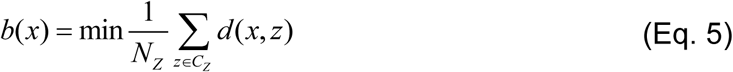

where *z* is a sample in *C_Z_*, and *N_Z_* is the total number of samples in *C_Z_*. The silhouette score of *x* is then defined as:

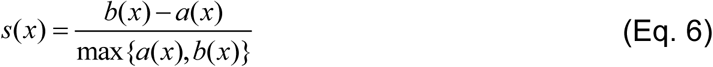

if the number of samples in *C_X_* is more than 1, otherwise *s*(*x*) = 0. We then computed the mean silhouette score over all samples.

### Decoder accuracy

To evaluate the performance of the decoders, we implemented a 4-fold cross validation, with training and testing sets each containing equal numbers of trials from all tasks. We used the coefficient of determination (R^2^), which indicates the proportion of variation of the actual EMGs that was predicted by the decoder. As EMGs are multi-dimensional, we computed a multivariate R^2^ (m-R^2^) in which, after computing the R^2^ for all the single dimensions, we found a weighted average across dimensions, with weights determined by the variance of each dimension. This was implemented using the “r2_score” function of the scikit-learn python package with “variance weighted” for the “multioutput” parameter.

## Results

### Global decoders failed to maintain prediction accuracy across tasks

We trained a series of Wiener filter-based decoders using data from each individual task, evaluated their accuracies within and across these tasks, and compared the results to those of global decoders trained on data encompassing all tasks of both environments. Figure 4A shows example EMG predictions for four representative muscles using the global linear decoder and decoders trained and tested exclusively on data from each task. The predictions by the within-task decoders tracked the actual EMGs well; however, the predictions by the global linear decoder were much worse. We further summarized the EMG decoding accuracy as m-R^2^ over all tests in Figure 4B. The single-task decoders achieved good EMG decoding accuracy for the trained task (entries along the diagonal), however, none predicted the EMGs accurately during different (off-diagonal) tasks. Furthermore, while performance of the linear global decoder (the second column from right) was well above the off-diagonal performance, it was dramatically lower than the decoders that were trained and tested on single tasks. We also tested a global LSTM decoder (the rightmost column in Figure 4B), which performed substantially better than the global linear decoder. For in-lab power and key grasp task, it performed better or equally well as the within-task linear decoders. Performance on the remaining tasks was marginally worse than the within-task linear decoders. These results are consistent with our illustration in Figure 1B: The within-task decoders achieved accurate linear fits for each cluster of datapoints, while the fits to each group of points by the global decoder were worse, as it endeavored to optimize the fit over all samples.

**Figure 4.**
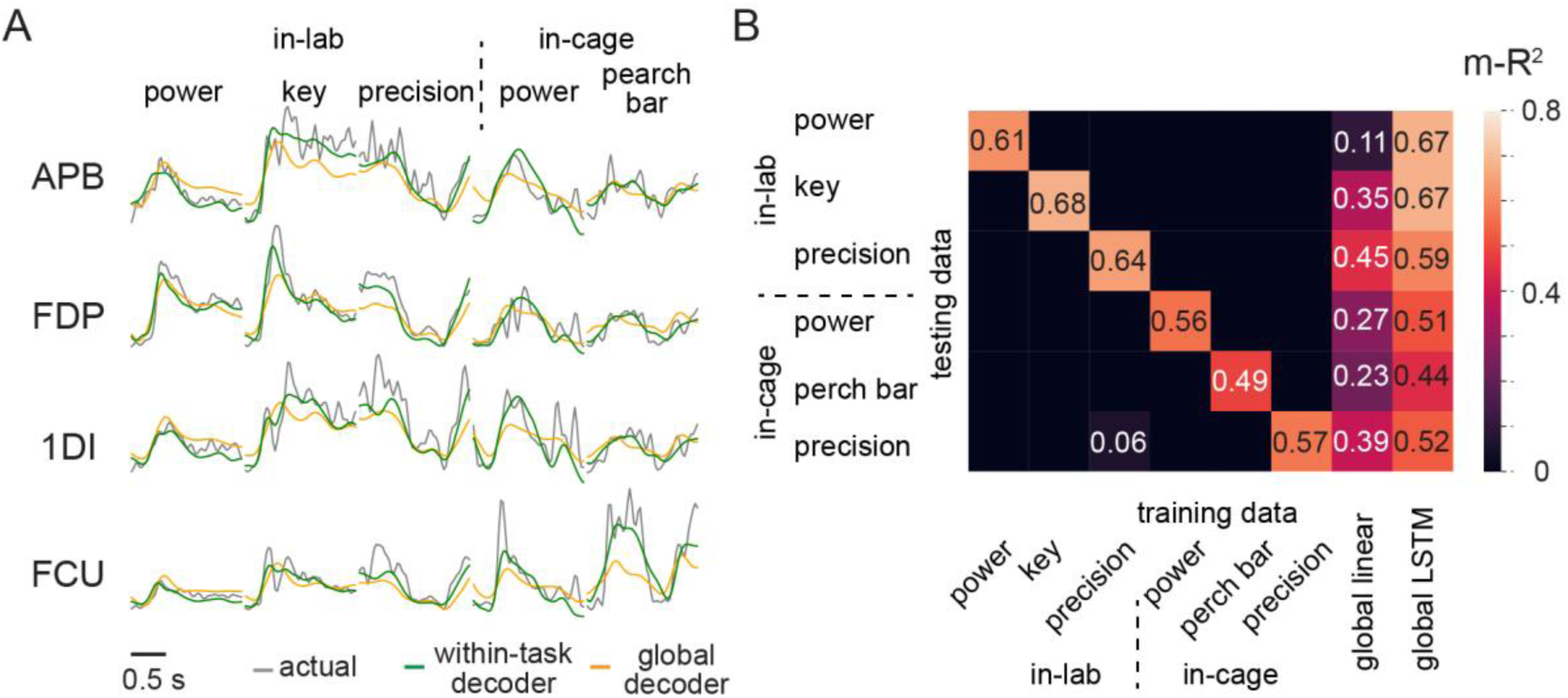
The performance of individual linear decoders and the linear global decoder over different tasks. **(A)** EMG prediction examples during one trial from each task when using both the within-task decoder and the linear global decoder fit with data from all tasks across the two environments. Each column shows a trial from one task and each row corresponds to a muscle. **(B)** A summary of EMG prediction accuracies using the global and single-task decoders. The prediction accuracy was measured by m-R^2^ across EMGs from 16 muscles. The m-R² values for the unlabeled (black) matrix locations are all negative.

### Task-related clusters emerge within the neural manifolds

We first determined the number of identifiable clusters in the neural manifolds following dimensionality reduction of the neural activity to 12 dimensions using either PCA or UMAP. We fit Gaussian mixture models with different numbers of clusters to these data and computed the silhouette scores over held-out testing trials. Figure 5A shows the silhouette scores for the PCA-based manifold, which remained consistently near 0.2, with no obvious peak. The low scores indicate that only rather indistinct clusters exist in the PCA-space, although the points are not distributed uniformly (which would yield score = 0). The absence of a prominent peak makes determining the number of clusters that best describe the clustering of the data difficult.

**Figure 5.**
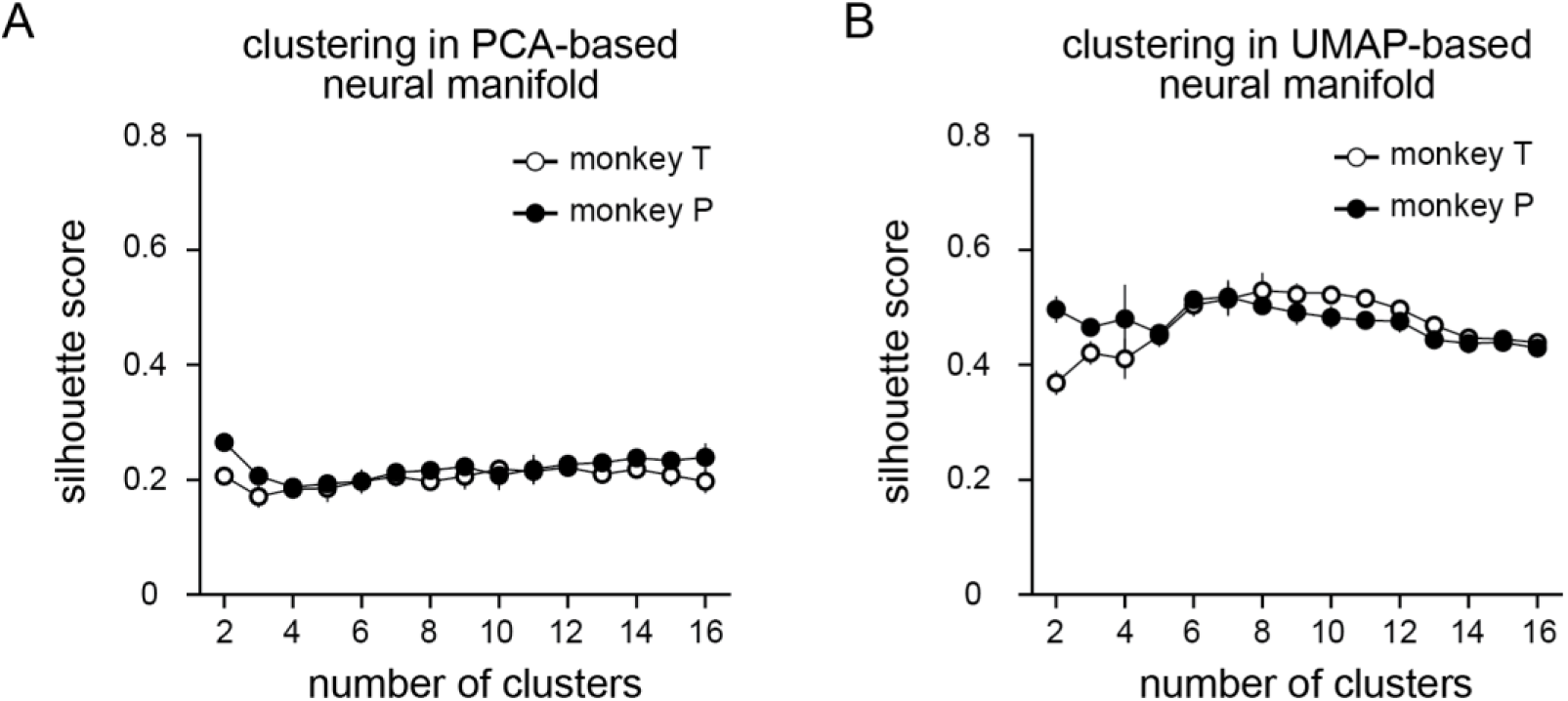
Determining the number of clusters in linear and nonlinear neural manifolds. (A) The silhouette scores indicate the quality of clusters found using Gaussian mixture models (GMMs), in this example, following dimensionality reduction by Principal Component Analysis (PCA) and **(B)** Uniform Manifold Approximation Projection (UMAP). The symbols represent mean silhouette values across 4 folds, and the error bars represent standard deviation.

The corresponding silhouette scores within the nonlinear UMAP-based manifold ranged from 0.4 to 0.6 (Figure 5B), more than twice that of the linear manifold. For monkey T, the highest score was achieved for eight clusters, although the peak was fairly broad. For monkey P, we observed similar results, with the highest score occurring for six or seven clusters, slightly greater than the number of recorded motor tasks. The more distinct clustering in the UMAP-space indicates that the actual neural manifold is significantly nonlinear.

We then examined the identified clusters from a representative session of monkey T to understand how they were associated with the tasks. Figure 6A shows the eight clusters of data samples identified by GMM in 3-dimensional PCA latent space. The data comprise equal numbers of trials for each of the six tasks, extracted from the continuously recorded data and concatenated. These clusters were adjacent in space, without clear boundaries, consistent with the silhouette score of 0.2 (Figure 5A). Figure 6B shows the time course of latent signals for 10 trials from each task, color-coded according to the cluster label of each data sample. It is notable that tasks of a similar nature, performed in different contexts, often shared clusters. For example, in-lab and in-cage power grasp shared clusters 7, 5, and 4, while precision grasp, both in the lab and cage, shared clusters 2 and 3. These plots also reveal cluster switching near certain behavioral events. For example, samples preceding force onset by up to 250 ms and corresponding to hand pre-shaping belonged largely to cluster 5 for both in-lab and in-cage power grasp. Samples immediately after force onset, corresponding to the maintained grasp, belonged to cluster 4. Similar observations could be made for the remaining tasks, but the transitions were less precisely aligned to task events. Since these tasks, including in-cage perch bar grasp, and both in-lab and in-cage precision grasp, were identified manually through video-based labeling, such variation may have been introduced by labeling deviations.

**Figure 6.**
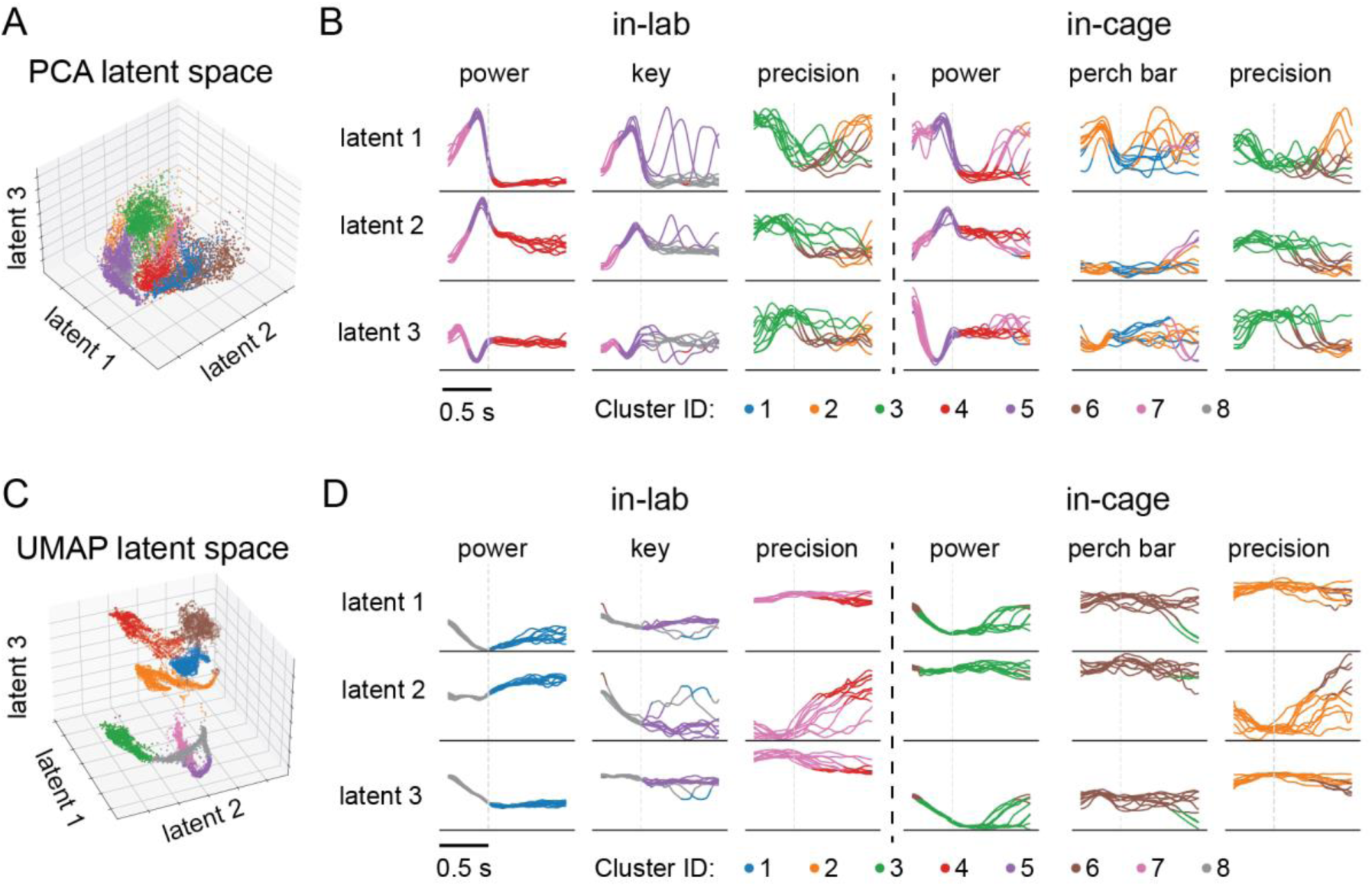
Spatiotemporal characteristics of neural manifold clusters. (**A**) Clustering (represented by different colors) of neural activity in linear neural manifold found by Principal Component Analysis (PCA). (**B**) Time plots of 10 representative individual trials for the first three dimensions in PCA space. The gray dashed line at 0.5 s in each sub-plot indicates the time of force onset during power and key grasp task, the moment when the monkey touched the food pellet during precision grasp, and the moment when the monkey touched the perch bar. (**C**) Clustering of neural activity in nonlinear neural manifold found by Uniform Manifold Approximation Projection (UMAP). Note that unlike PCA, the latent components of UMAP were not ordered by variance. (**D**) Corresponding time plots of the first three UMAP dimensions.

Figure 6 C, D shows the analogous clustering in the nonlinear UMAP space. In contrast to the linear clusters, these were clearly separated. Each of the in-cage tasks and the in-lab precision grasp task had exclusive clusters (7: in-lab precision grasp, 3: in-cage power grasp, 6: in-cage perch bar grasp, 2: in-cage precision grasp). In-lab power grasp and key grasp shared cluster 8 before force onset, then separated into clusters 1 and 5 after force onset. Notably, when GMM was set to identify more than eight clusters in the UMAP space, the additional clusters were quite small, presumably reflecting occasional, atypical behaviors, while the correspondence between well-defined tasks and the primary clusters persisted. We observed similar results in sessions from monkey P. Together, these observations suggest that neural activity in both linear and nonlinear low-dimensional manifolds form identifiable clusters. These clusters are readily interpretable given their correspondence to either the category or sub-phase of the tasks.

### Piecewise linear decoding outperformed both linear and nonlinear global decoders

We evaluated the performance of the piecewise linear decoders with various numbers of clusters and compared it to that of both linear and nonlinear global decoders. Figure 7 illustrates these results from a representative session for each monkey. We calculated the average m-R² between the actual and predicted EMGs on three experimental sessions for each monkey using 4-fold cross-validation on testing data that each included all tasks. The piecewise linear decoders significantly outperformed the global linear decoders in accuracy, with just two clusters, for both PCA and UMAP clustering (monkey T, m-R^2^ for piecewise linear decoders: 0.57 ± 0.02 for PCA, 0.62 ± 0.01 for UMAP (mean ± s.e.); m-R^2^ for global linear decoder: 0.48 ± 0.01; both paired t-test comparisons were p<0.001. monkey P, m-R^2^ for piecewise linear decoders: 0.57 ± 0.02 for PCA, 0.59 ± 0.01 for UMAP; m-R^2^ for global linear decoder: 0.53 ± 0.01; both paired t-test comparisons were p<0.001). Their performance continued to improve and plateaued at 14 clusters for monkey T (no significant difference between 14 and 16 clusters; p = 0.16, paired t-test), and at 13 for monkey P (p = 0.06, paired t-test between 13 and 15 clusters) for PCA piecewise linear decoders. In the PCA-based scenario, m-R^2^ for the piecewise linear decoders significantly surpassed even the global LSTM decoder at 10 clusters for monkey T (0.69 ± 0.01, mean ± s.e. and 0.68 ± 0.01, respectively; p = 0.02, paired t-test) and 12 clusters for monkey P (0.64 ± 0.01 and 0.63 ± 0.01, respectively; p = 0.008, paired t-test). For the UMAP-based clusters, this superior performance occurred already with only 6 clusters for monkey T (0.69 ± 0.01, mean ± s.e. and 0.68 ± 0.01, respectively; p = 0.03, paired t-test) and 10 clusters for monkey P (0.64 ± 0.01, mean ± s.e. and 0.63 ± 0.01, respectively; p = 0.01, paired t-test).

**Figure 7.**
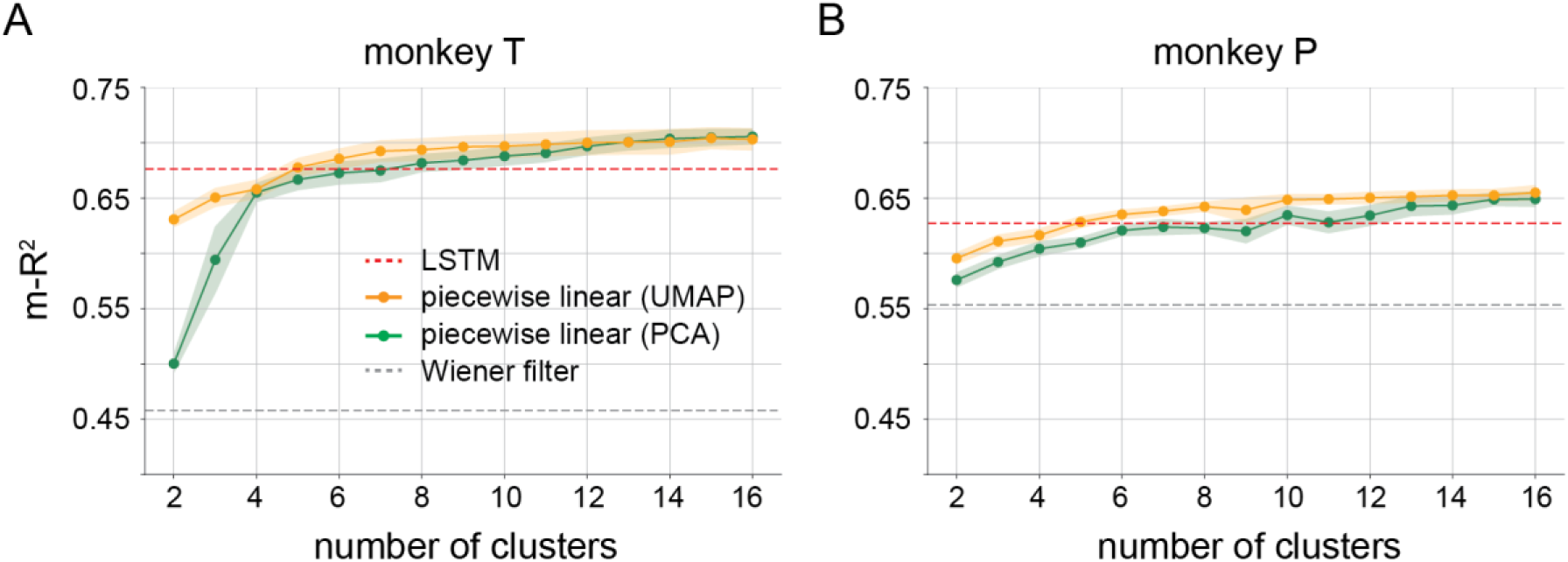
The EMG prediction accuracies for the global decoders and the piecewise linear decoders, tested with different numbers of clusters. For each type of global decoder (LSTM or Wiener filter) there is only one m-R^2^ value, computed across 16 EMG channels on a testing set that included trials from all in-lab and in-cage tasks.

Performance also differed between the PCA and UMAP spaces with relatively few clusters. UMAP outperformed PCA with fewer than 12 clusters for monkey T (p < 0.001 for 12 clusters or less except p = 0.003 for 4 clusters, p > 0.05 for all clusters above 12, paired t-test), and 8 for monkey P (p < 0.001 for 7 clusters or less except p = 0.03 for 4 clusters, p = 0.04 for 8 clusters, p > 0.05 elsewhere).

We further compared the performances of the decoders on individual tasks (Figure 8). For each monkey and pair of decoders, we calculated the difference in m-R^2^ prediction accuracy for individual tasks across three sessions. We first compared the performance of PCA-based cluster to that of UMAP (Figure 8A). Datapoints below zero indicate that UMAP had overall better performance compared to PCA. However, the performance advantage was modest, generally not exceeding 0.1, and came at a cost of significant computational overhead when computing the nonlinear projection compared to PCA. Given this tradeoff, a PCA-based piecewise linear decoder may be a more feasible solution for practical iBCI systems, therefore, we only tested PCA-based piecewise linear decoders in further analyses.

**Figure 8.**
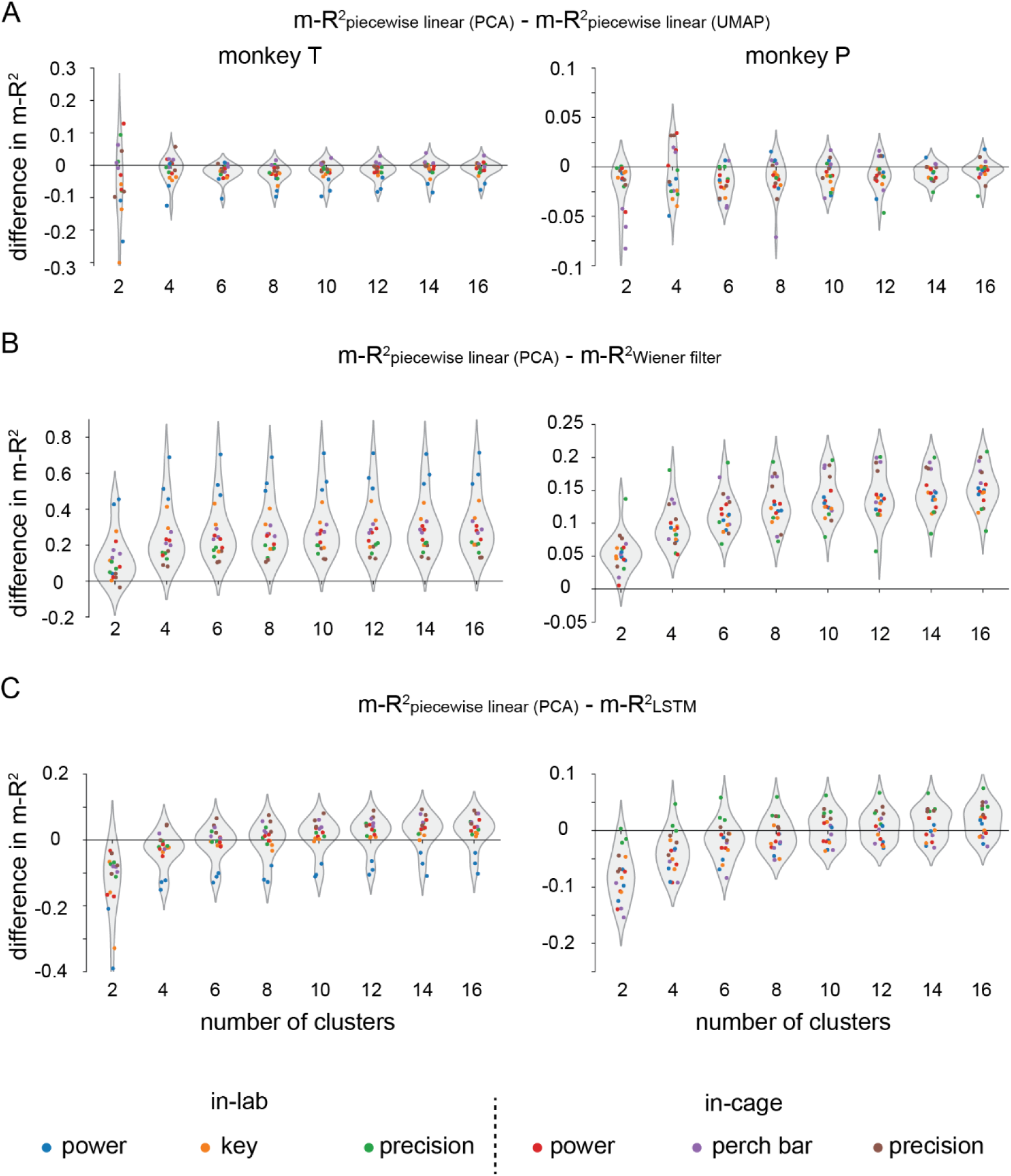
Comparisons between different types of decoders for each task. **(A)** The difference in decoding accuracy between piecewise linear decoding in PCA-space and UMAP-space. Each dot shows the comparison for one task in one (of three) dataset. A value above zero means piecewise decoding had superior performance. **(B)** Same format, comparing piecewise PCA and a global linear decoder. **(C)** Same format, comparing piecewise PCA and a global LSTM decoder.

For all numbers of clusters greater than one (where the approaches are equivalent) the PCA piecewise linear decoders consistently outperformed the global linear decoders (Figure 8B). With two clusters, the global LSTM decoder performed better for all tasks, but it gradually lost this advantage as the number of clusters increased. For monkey T, the piecewise linear decoders surpassed the global LSTM decoder for all but the in-lab power grasp task with 10 clusters. For monkey P, the piecewise linear decoder achieved better decoding accuracy with 12 clusters for most in-cage tasks, but the in-lab power and key grasp tasks remained slightly less accurate, even with 16 clusters (Figure 8C).

### Piecewise linear decoders yield accurate decoding even from continuously recorded data

In real-world iBCI applications, there is usually no predefined trial structure or fixed set of motor behaviors. Consequently, the decoder must work continuously to produce accurate predictions once trained. To assess whether the piecewise decoders could meet this requirement, we trained them using data from the initial 15 minutes of an in-cage recording session without selecting individual trials or identifying specific types of tasks. Subsequently, we evaluated their performance on the remaining continuously recorded data.

As noted above, the slight accuracy advantage of UMAP-based piecewise decoding (Figure 8A) is computationally expensive. Since the number of data samples in continuous recordings was ∼10 times greater than in the trial-concatenated case, training the neural networks required to compute the parametric UMAP projection for 15 minutes of continuous data demanded >1 hour of CPU time, as did performing inference on the remaining 30 minutes of data. Although a GPU could accelerate the calculations, the improvement in time efficiency was modest. Consequently, it was feasible only to do this analysis in linear space. Figure 9 shows the results from one representative session for each monkey. Because of the greater complexity and more widely varied behaviors of these datasets, we set the number of clusters to 12. We computed m-R^2^ in one-minute windows and used linear fits to characterize decoding accuracy throughout the session as a function of time. For both monkeys, the piecewise decoding performance was better than both global linear and LSTM decoders (Figure 9A, B). Although predictions were somewhat less accurate than for the trial-concatenated data (Figure 7), the small difference was surprising, considering that the continuous data had nearly an order of magnitude more datapoints representing less well-defined defined behaviors. It had been well acknowledged that the drifting of neural recordings may affect the stability of iBCI decoding accuracy [33], even within the same session [34]. We observed slow declines in m-R^2^ over the 30-minute testing data, characterized by the negative slopes of the linear fits for all decoders, which suggests the existence of such instability. The slopes for the piecewise linear decoders were smaller than those of the LSTM decoder. Figure 9 C, D show segments of continuously recorded EMGs and the corresponding piecewise linear predictions for monkey T and monkey P, respectively. It was notable that even though the EMGs were more variable compared to the trial-based case (Figure 4B), the piecewise linear decoders made accurate predictions by switching between the appropriate decoders. Together, these results reinforce the feasibility of using piecewise linear decoding in practical iBCI use, especially for an FES BCI, which requires continuous EMG predictions to modulate the electrical stimulator.

**Figure 9.**
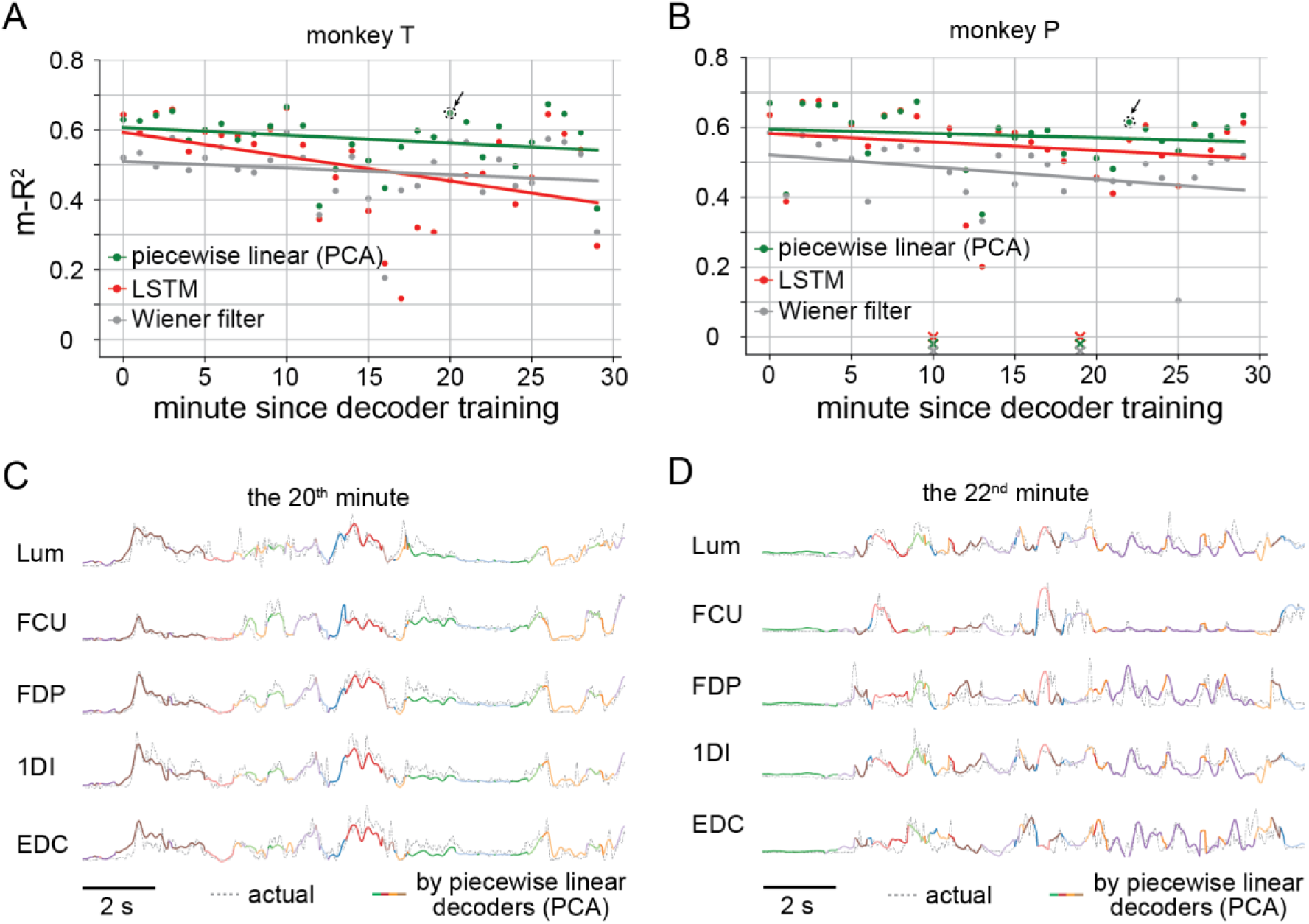
EMGs decoded from continuously recorded M1 signals as monkeys moved freely in the cage. **(A)** The accuracies of three types of decoders for data from monkey T. Each dot represents decoding accuracy for 16 muscles (m-R^2^) computed in one-minute windows with a particular decoder (indicated by color). **(B)** Corresponding results for monkey P. **(C)** A segment of actual EMGs and the predictions using piecewise linear decoders from the session of monkey T in (A). This segment was from the 20^th^ minute since decoder training. The colors in the predictions indicate the cluster of the neural signals belonged to at the current time step. **(D)** EMG samples and predictions from the session of monkey P in (B), which was from the 22^nd^ minute since decoder training.

## Discussion

Most iBCIs have been tested in a relatively limited range of motor behaviors, and virtually all effect purely kinematic control. The FES BCI we developed previously [12] has the advantage that users can control both postures and forces with the same interface, but the ever-changing control dynamics present a challenge to the decoder. A user’s experience with the iBCI would be significantly impacted by any inability to work effectively across tasks [35]. Here, we have introduced a novel approach to enable iBCIs to handle multiple tasks and contexts, specifically for an EMG decoder, but a similar approach should work for decoders of movement kinematics [8,36,37] or grasp force [9,38]. In the interest of developing highly generalizable decoders, there have been recent exciting proposals to develop “large cortical models” that mimic the training of the remarkably successful large language models [21], thereby to learn huge dictionaries of motor cortical dynamics [22,23]. However, these approaches require vast amounts of data and intensive GPU-based training; our approach requires neither. We developed a pipeline that finds clusters of neural activity within low-dimensional manifolds and then trains a linear decoder to predict EMGs for each cluster. This allowed us to approximate the complex nonlinearities [39,40] in neural activity in a piecewise linear manner. We compared this piecewise linear decoding with global Wiener filter and LSTM-based decoding, both trained on data from all tasks. Surprisingly, the piecewise linear approach slightly outperformed even the LSTM.

### Challenges for building iBCI decoders to handle multiple tasks

iBCI systems typically enable users to handle one type of task in a fixed setting, for example, moving several fingers [41], a 2D cursor [6], a virtual arm avatar [42], or an anthropomorphic robot arm [8]. While a linear decoder can effectively represent the relationship between M1 activity and motor outputs for a single task, for a broader range of tasks, the effect of nonlinearities in the mapping becomes more pronounced. Indeed, in the current study a linear EMG decoder, trained with data from all the tasks, performed relatively poorly on each component task (Figure 4B). Similar observations have been made for wrist muscles under different loading conditions, and for decoding the velocity of both initial and corrective movements during reaching [14]. These results highlight the impact of the nonlinearity between M1 activity and motor outputs for iBCI systems. Such nonlinearities may originate from downstream processes such as gain mechanisms in subcortical structures or the spinal cord [20,43,44], or from cognitive or sensory feedback inputs to M1 [45]. It is typically not feasible to model these nonlinearities explicitly. If we instead parametrize them using artificial neural networks, such as LSTM-based decoders, we can achieve significantly higher performance (Figure 4B). However, selecting optimal parameters for these decoders during training can be difficult, yet may make the difference between mediocre and excellent performance [46].

Instead, we hypothesized that we could approximate the nonlinearities across tasks using a set of linear decoders trained on samples from clusters occurring naturally in low-dimensional neural manifolds. Our approach tests the hypothesis (illustrated in Figure 1B), that simple linear decoders can be used to make accurate predictions of EMG throughout a broad range of contexts and motor behaviors, if they can approximate the complex, nonlinear geometry of the neural landscape in M1. Remarkably, not only was our piecewise linear decoder more accurate than a global linear one, but it even sightly outperformed a global LSTM decoder, making it a viable option for multi-task iBCI systems.

The most salient factor determining the accuracy of our piecewise decoding approach is the number of clusters. Although the analysis of silhouette scores alone did not offer a definitive means to select an optimal number of clusters (Figure 5), there were significant changes in decoding accuracy as the number of clusters increased (Figure 7). Piecewise decoders began to outperform the global nonlinear decoder when using eight clusters, just above the number of identified tasks. However, as the number of clusters increased further, performance saturated, in part because the “additional” clusters had very few data samples so the corresponding decoders could not be adequately trained. Another factor that influenced performance is the dimensionality reduction method used to identify the neural manifold. As shown in Figure 7, decoders trained on clusters within the UMAP latent space had a small but consistent advantage that vanished when the number of clusters exceeded 12. For this reason and given the reduced computational complexity of the linear space, we concluded that piecewise decoders based on clusters within the PCA latent space are a more efficient solution for practical iBCI use.

### Organization of the neural activity in the low-dimensional manifold

Previous studies in species ranging from drosophila, to mice, to monkeys, have used dimensionality reduction to process movement kinematics and identify clusters within a low-dimensional behavioral subspace [47–49]. This allows complex behaviors to be decomposed into simpler sub-behaviors, corresponding to the identified clusters, which have been called “behavioral motifs”. It is reasonable to assume that there may be similar “neural motifs” controlling these behaviors. These neural motifs may be found at different locations in the neural manifold, with identifiable topological relationships between each other, revealing how neural activity is organized during motor control. However, verifying this hypothesis has been challenging, due to the limited variety of motor behaviors for which neural recordings were available. This is especially true for studies with monkeys, as they generally need to be restrained in laboratory setting and connected to the equipment needed to record neural signals. Our group previously examined the neural activity recorded from monkeys in the lab during several types of constrained wrist and grasp tasks and found the underlying (linear) neural manifolds all had similar orientation [50].

Our use of wireless recordings [51–53] within a cage allowed us to expand the range of tasks explore how the neural activity underlying these tasks in very different contexts were organized in the manifold. We recently computed the dimensionality and linearity of neural activity in these two settings using methods developed and validated using synthetic data [54,55]. We found the in-cage task dimensionality was approximately 50% higher than those in the lab, and that the in-cage manifolds tended to be more nonlinear. By studying the clustering of neural activity in both linear and nonlinear spaces, we were able to infer even more details about the organization of neural manifolds, such as how the sub-regions of the manifold are related to the tasks, and the topological relationships between them. Unlike nonlinear space, clusters within the linear manifolds, were unexpectedly related to sub-phases of tasks. For instance, hand pre-shaping clusters were commonly identified in both in-lab power and key grasp, as well as in in-cage power grasp (Figure 6A and B). The nonlinear view of the neural manifold, while more challenging computationally, may better approximate its true curved geometry [39,54,56], The more clearly-separated, task specific clusters (Figure 6 C and D) suggest that neural activity for different tasks and contexts may reside in different regions of a curved nonlinear manifold. Thus, in addition to its success at decoding complex movements, our approach can enhance our understanding of the organization of M1, providing an intuitive view of the complex processes underlying multiple motor tasks and across different contexts.

### Limitation and future directions

In summary, we have tested a piecewise linear approach to EMG decoding in a multi-task iBCI that is easy to implement, yet as accurate as a nonlinear RNN. While promising, we recognize several limitations in the current study. Although our approach can produce accurate EMG predictions from data that span multiple tasks, it requires training data from all tasks for good performance. We performed leave-one-task-out tests using piecewise linear filters, global Wiener filters, and LSTM decoding. All failed to predict the motor outputs of the left-out task. Extrapolation beyond the well-explored region of a nonlinear manifold is always likely to fail. However, by further studying the topology of the curved manifolds and the geometric relationships between the clusters, we may be able to build decoders through interpolation across the curved manifold, at least for novel tasks having activity that falls within the convex hull of the training data.

## Acknowledgments

We thank Ann Kennedy and Ege Altan for valuable discussions. We also thank former members of the Miller Limb Lab, including Robert H. Powell, Juliet Heye and Qiwei Dong, for their contributions to data collection and set-up of experimental equipment. The work was supported by grants to L.E.M. (R01 NS053603, R01 NS074044).

## Author contributions

X.M. and L.E.M. designed research; X.M and K.L.B., performed data recording and experiments; X.M. and F.R. analyzed data; X.M, F.R., and L.E.M. wrote the paper.

